# Ligand Screening and Discovery using Cocktail Soaking and Automated MicroED

**DOI:** 10.1101/2025.02.18.638921

**Authors:** Jieye Lin, Marc J. Gallenito, Johan Hattne, Tamir Gonen

**Affiliations:** Department of Biological Chemistry, University of California, Los Angeles, 615 Charles E. Young Drive South, Los Angeles, California 90095, United States; Department of Physiology, University of California, Los Angeles, 615 Charles E. Young Drive South, Los Angeles, California 90095, United States; Howard Hughes Medical Institute, University of California, Los Angeles, Los Angeles, California 90095, United States

## Abstract

Cocktail soaking using single-crystal X-ray diffraction (SC-XRD) has previously allowed high-throughput crystallographic screening of ligands against protein targets. However, protein microcrystals are not amenable to this approach if they are too small to yield strong diffraction patterns. In this study, we developed a workflow integrating cocktail soaking with automated microcrystal electron diffraction (MicroED) to allow rapid ligand screening, structure determination, and binding analysis directly from microcrystals. This can improve the successful hit rate, because binding is often more efficient when smaller crystals are soaked in the ligand. The approach was validated with known ligands of thermolysin and identified novel binding interactions for ligands of proteinase K. The structures of multiple protein-ligand complexes, including ligands with weak binding affinities, could be solved quickly. Their estimated relative binding affinities are in good agreement with previous work and independent microscale thermophoresis (MST) measurements.

## 1 Introduction

Macromolecular crystallography remains the leading technique for 3D structural determination of protein-ligand complexes at atomic resolution, because the ligand and its interactions with the protein can be determined from density maps.^1^ Crystallographic ligand screening was traditionally limited by low throughput until 1997, when Verlinde *et al*. introduced the concept of “cocktail soaking”, which significantly improved the ligand screening rates. After soaking triose-phosphate isomerase (a glycolytic enzyme) in four cocktails of 128 anti-trypanosomiasis drugs, they ultimately identified one binding hit by analysis of Fo-Fc maps.^2^ The method has since been widely applied in drug discovery targeting various disease-related proteins like urokinase (a kinase highly expressed in invasive cancers)^3^ and nucleoside 2-deoxyribosyltransferase (related to Chaga’s disease)^4^. Additionally, cocktail soaking has revealed allosteric binding sites and unknown functions on target proteins.^5,6^ The efficiency of cocktail screening has been significantly enhanced by advances in robot-assisted soaking^7,8^ and automated single-crystal X-ray diffraction (SC-XRD) data collection.^8-10^

Achieving high occupancy of ligand in crystals typically requires highly concentrated ligands and long incubation time^11^. The process also risks cracking or dissolving the protein crystals, which compromises the diffraction quality.^11^ On the contrary, soaking ligands into microcrystals is generally more effective, as their smaller crystal volumes facilitate ligand diffusion.^12,13^ However, most microcrystals are unsuitable for conventional synchrotron-based X-ray diffraction due to their small sizes (<5 µm),^14^ and access to X-ray free-electron laser (XFEL) facilities remains globally limited.^15^ These shortcomings can efficiently be addressed with microcrystal electron diffraction (MicroED).

MicroED is well-suited for 3D structural analysis, requiring crystals only a billionth size of those needed for conventional X-ray diffraction.^16-18^ MicroED has been widely adopted for elucidating the structures of pharmaceutical small molecules,^19,20^ proteins,^21,22^ and protein-ligand complexes.^23^ Automated MicroED has significantly increased throughput, such that hundreds or thousands of datasets are sequentially collected grain-by-grain, enabling efficient structural and compositional analysis of mixtures^24,25^ (Figure 1). MicroED is a promising method for cocktail screening, as indicated in our prior study, where the binding of 5-amino-2,4,6-triodoisophthalic acid soaked into proteinase K, was determined with superior efficiency and occupancy compared to larger crystals under the same condition.

**Figure 1.**
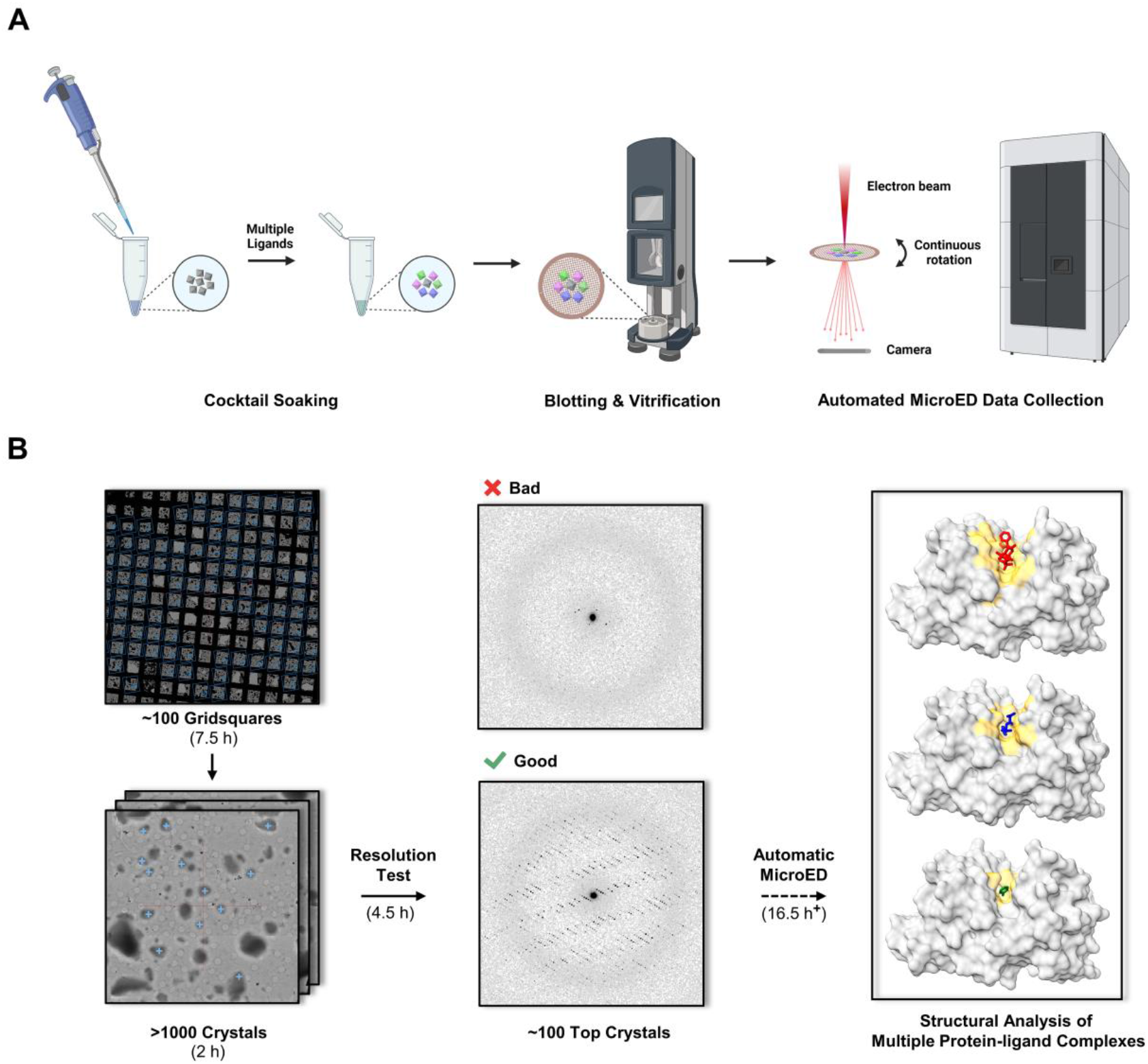
The workflow of (A) cocktail soaking and (B) automated MicroED data collection. Typical data collection was performed in two major steps: (1) a pre-resolution test of over 1000 microcrystals in less than 5 hours to estimate data quality; and (2) automated MicroED data collection of the top 100 crystals overnight.

In this study, we used automated MicroED to rapidly screen for ligand binding in protein microcrystals soaked in ligand cocktails (Figure 1).^24,25^ *Multiple complexes were determined for the same protein and the relative binding affinities of each ligand were assessed*. We first validated the workflow using three known ligands of thermolysin (**TLN**): phosphoramidon disodium salt (ligand **1**), thiorphan (ligand **2**) and aniline (ligand **3**).^26-28^ We then extended this workflow to uncover the unknown protein-ligand bindings of three potent ligands for proteinase K (**PK**): benzenesulfonyl fluoride hydrochloride (ligand **4**), diisopropyl fluorophosphate (ligand **5**) and a tetrapeptidyl chloromethyl ketone compound, MeOSuc-Ala-Ala-Pro-Phe-CH_2_Cl (ligand **6**). Their resulting refined structures are consistent with previous studies (Figures 2 and 3)^29,30^ and our estimated relative binding affinities agree with independent microscale thermophoresis (MST) measurements (Figure 4). This study demonstrates that cocktail soaking and MicroED can provide a robust means for screening ligands against protein targets.

**Figure 2.**
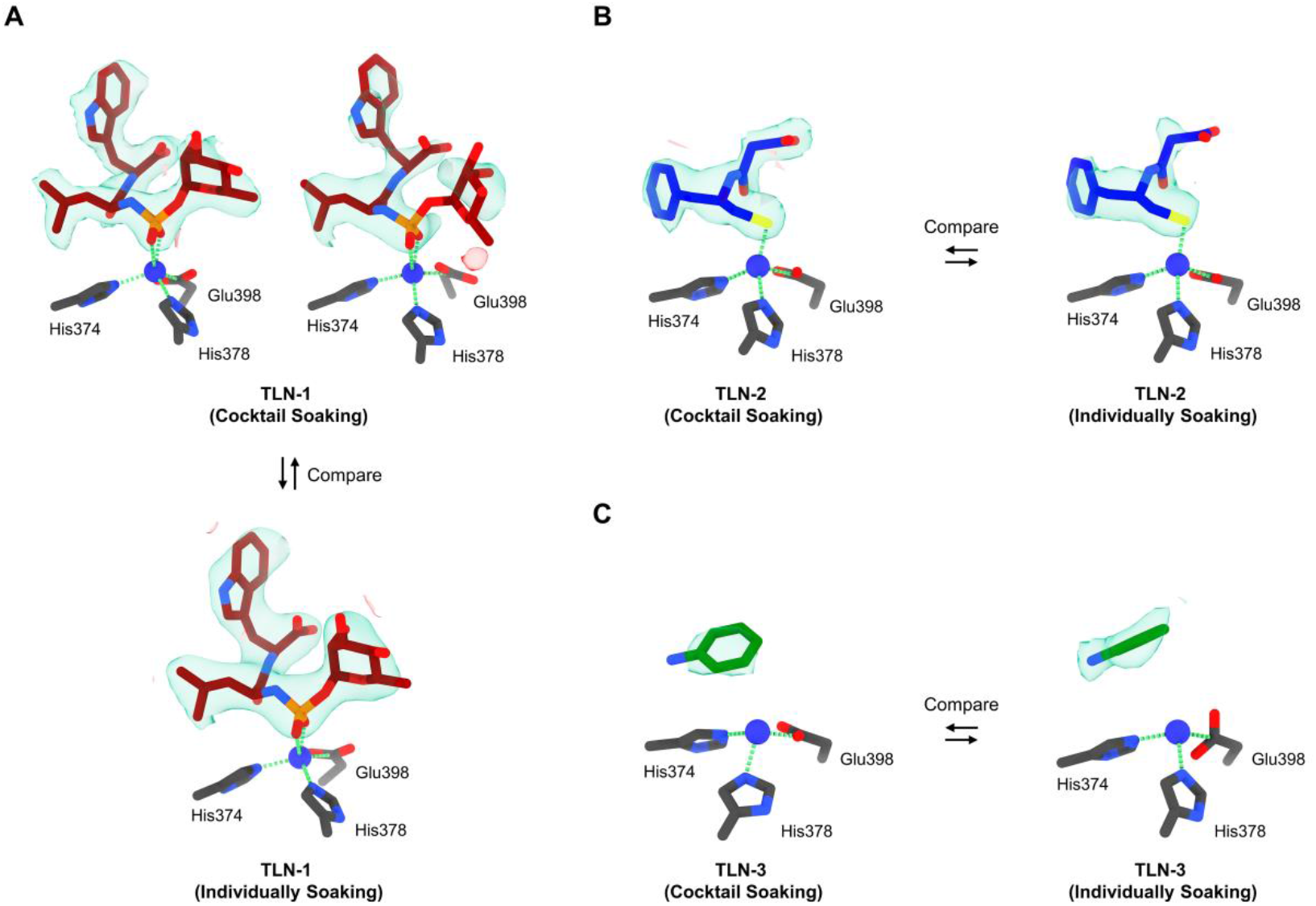
Comparison of the structure and charge densities of thermolysin (**TLN**) in complex with ligands **1**-**3** using two soaking approaches. (A) Top-left, **TLN**-**1** from cocktail soaking set A1 (Fo-Fc map: 2.5σ contour); Top-right, **TLN**-**1** from cocktail soaking set A2 (Fo-Fc map: 2.1σ contour); Bottom, **TLN**-**1** from individual soaking set B (Fo-Fc map: 2.5σ contour). (B) Left, **TLN**-**2** from cocktail soaking set A2 (Fo-Fc map: 2.5σ contour); Right, **TLN**-**2** from individual soaking set C (Fo-Fc map: 2.5σ contour). (C) Left, **TLN**-**3** from cocktail soaking set A2 (Fo-Fc map: 2.5σ contour); Right, **TLN**-**3** from individual soaking set D (Fo-Fc map: 3.0σ contour). The carbon atoms for ligands **1**-**3** were colored in red, blue and green, respectively. Ligands **1**-**3** densities (Fo-Fc) were colored in green and red. Thermolysin was abbreviated as “**TLN**”.

**Figure 3.**
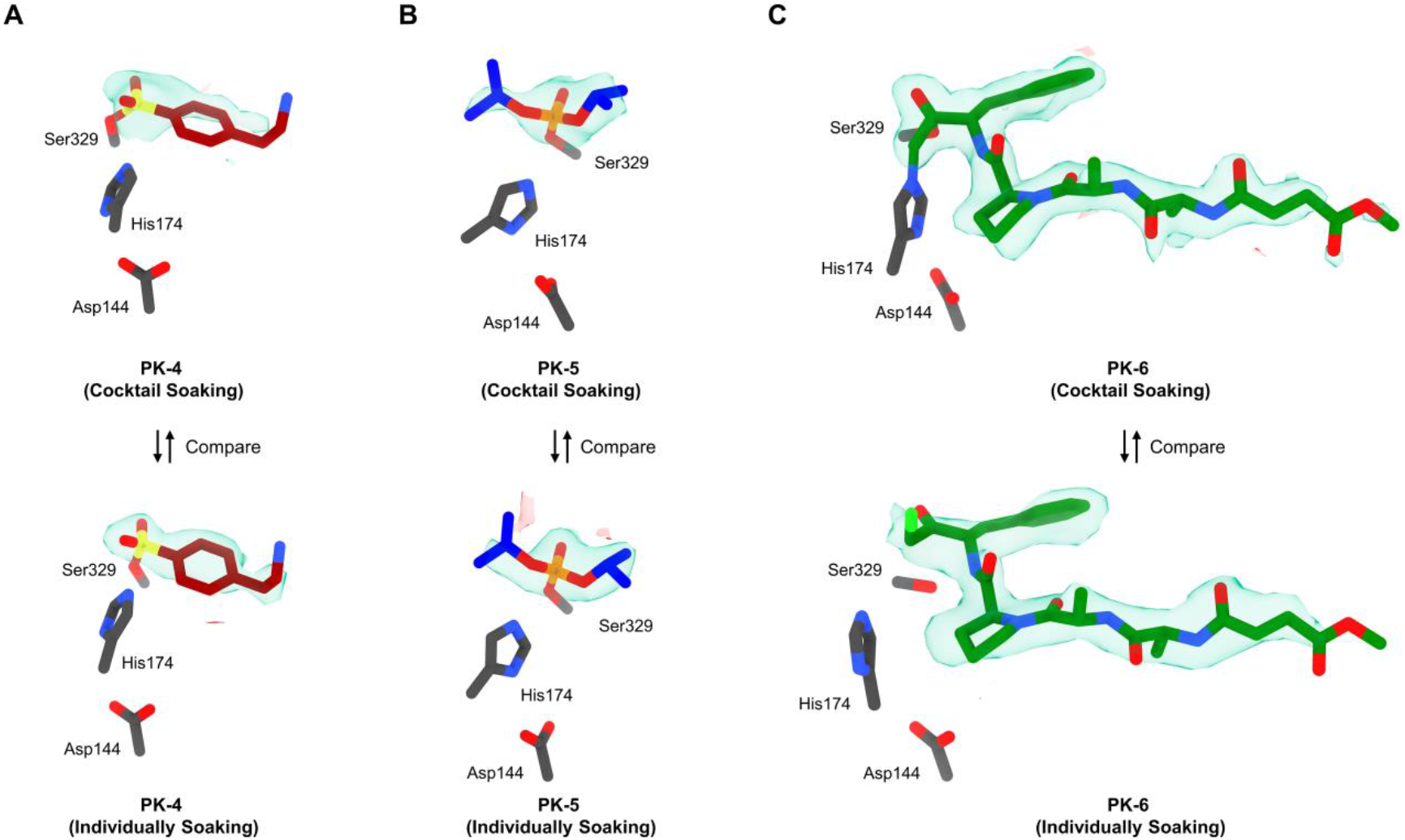
Comparison of the structure and charge densities of proteinase K (**PK**) in complex with ligands **4**-**6** using two soaking approaches. (A) Top, **PK**-**4** from cocktail soaking set E (Fo-Fc map: 2.3σ contour); Bottom, **PK**-**4** from individual soaking set F (Fo-Fc map: 2.3σ contour). (B) Top, **PK**-**5** from cocktail soaking set E (Fo-Fc map: 2.5σ contour); Bottom, **PK**-**5** from individual soaking set G (Fo-Fc map: 2.5σ contour). (C) Top, **PK**-**6** from cocktail soaking set E (Fo-Fc map: 2.5σ contour); Bottom, **PK**-**6** from individual soaking set H (Fo-Fc map: 2.5σ contour). The carbon atoms for ligands **4**-**6** were colored in red, blue and green, respectively. Ligands **4**-**6** densities (Fo-Fc) were colored in green and red. Proteinase K was abbreviated as “**PK**”.

**Figure 4.**
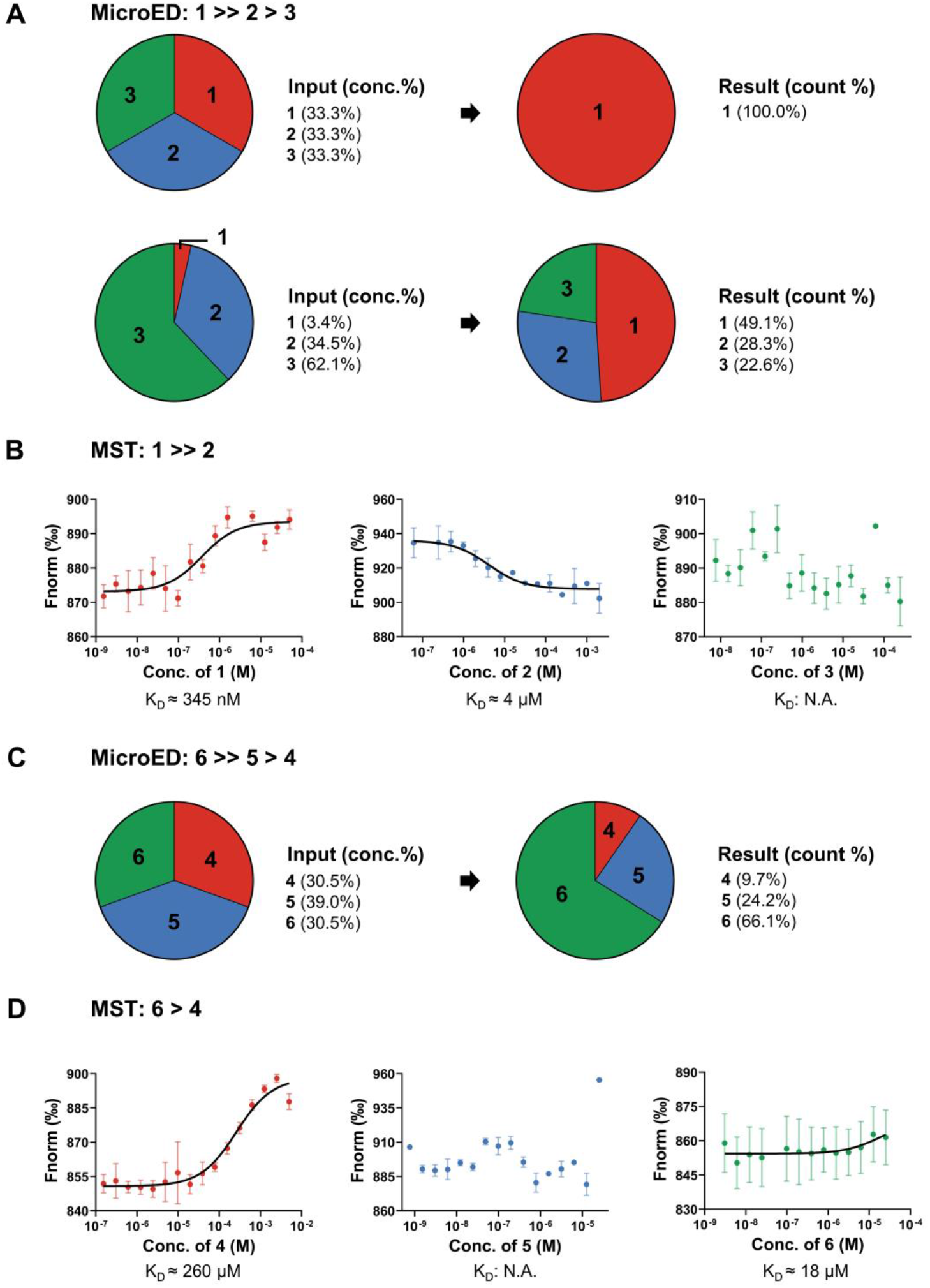
Relative binding affinities of ligands **1**-**6** determined by MicroED and MST. (A) Relative binding affinities determined by MicroED after analysis of **TLN**/**1**-**3** complexes in cocktail soaking sets A1-A2. (B) K_D_ values of ligands **1**-**3** measured by MST. (C) Relative binding affinities determined by MicroED after analysis of **PK**/**4**-**6** complexes in cocktail soaking set E. (D) K_D_ values of ligands **4**-**6** measured by MST. K_D_ was calculated by Equation 1 (replicate 2 times).

## 2 Results

### 2.1 Method Validation: Thermolysin in Complex with Ligands 1-3

Thermolysin (**TLN)** is a protein originally isolated from *Bacillus thermoproteolyticus*,^31^ containing a Zn ion coordinated by His374, Glu398 and His478 residues in its active sites (Figure S2 in Supporting Information).^32,33^ **TLN** can form complexes with three known ligands **1**-**3** (PDB entries: 1TLP, 1Z9G, 3MS3)^26-28^ in the same binding site. Soaking **TLN** in a cocktail of ligands **1**-**3** is expected to yield crystals in which the active sites are occupied according to the binding affinities (K_D_) and concentrations of the respective ligands, since all the ligands bind non-covalently to the same site.

In set A1, thermolysin crystals were soaked for 5 min in a cocktail of equal concentrations of the three ligands **1**-**3** (Table 1). Three individual soaking sets B-D only contain a single ligand were set as references to demonstrate the ligand densities (Table 1). Data from 115 crystals were rapidly collected by automated MicroED overnight (Figure 1B) and after trimming (see “Method” for details) 65 structures were solved by molecular replacement without merging.^34^ The shape of Fo-Fc densities in the binding site was overall well-defined for all structures (literature^26^ and reference set B, Figure 2A), but only fit ligand **1** indicating that its binding affinity is much greater than those of ligands **2** and **3** (Figure S3 in Supporting Information).

**Table 1.**
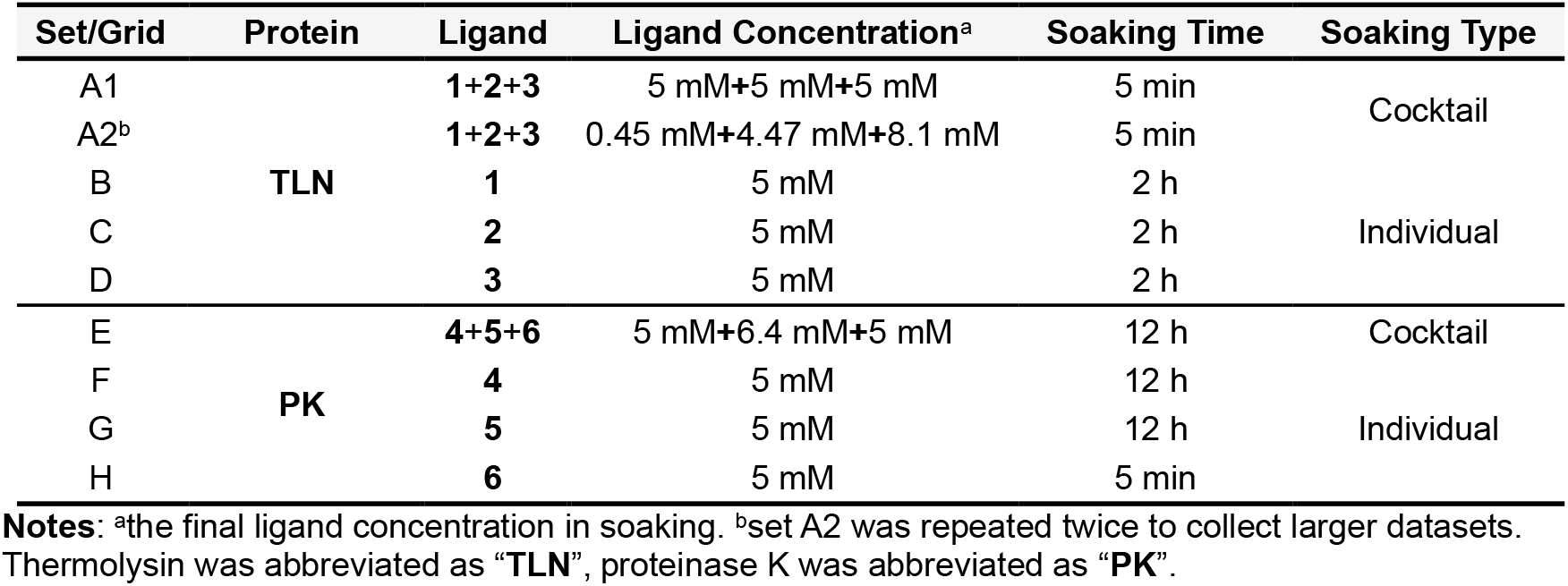
Soaking setups for Thermolysin and Proteinase K.

In a second cocktail (set A2), ligands **1**-**3** were mixed in approximately 1:10:20 ratios (Table 1). The concentrations were set corresponding to the binding affinities calculated by AutoDock Vina,^35,36^ *i*.*e*. -12.3 kcal/mol for **TLN**-**1**, -6.1 kcal/mol for **TLN**-**2**, and -4.3 kcal/mol for **TLN**-**3**. A total of 102 structures were solved (Figure S4 in Supporting Information) from 225 datasets, of which, 53 structures were found to exhibit Fo-Fc density in their binding sites (Figures 2 and 4A). The data quality in set A2 was lower than in set A1, likely due to the presence of crystals binding different ligands (reference sets B-D, Figure 2). Despite these limitations, the densities retained distinct ligand density shapes (Figure 2). For example, in maps from crystals where ligand **1** was predominantly bound, the tryptophan is clearly visible but the density for the L-ramnopyranoside moiety is weak (Figure 2A), possibly due to an increased flexibility or competition from ligands **2** and **3** (Figure S6A in Supporting Information). The density of ligand **2** in set A2 showed better occupancy than previously observed^27^ (reference set C, Figure 2B). The density of ligand **3** in sets A2 and D was generally complete and showed high consistency (Figure 2C). The terminal amine was visible in set D but missing in set A2, likely due to their resolution differences (Figure S6C in Supporting Information). Ligand **2** showed moderate binding affinity, as more **TLN**-**2** complexes were found in set A2 despite using half the soaking concentration of ligand **3**. As no conserved binding geometry was observed for **TLN**-**3**, it is possible that it resulted from non-specific diffusion.^28^ MST measurements of K_D_ values for ligands **1**-**3** were 345 nM for ligand **1**, 4 µM for ligand **2**, and N.A. for ligand **3**, showing consistency with literature^29,30^ and MicroED results (Figures 4A-B).

### 2.2 Unknown Protein-ligand Binding: Proteinase K in Complex with Ligands 4-6

Proteinase K (**PK**) is a serine proteinase originally purified from *Tritirachium album*,^37^ featuring an active site with oxyanion triad composed of Asp144, His174 and Ser329 residues (Figure S2 in Supporting Information).^33^ **PK** is known to be inhibited by many ligands but lack published structures in literature, for example, ligand **4** can form an O−S bond with Ser329 residue using the same mechanism as phenylmethylsulfonyl fluoride (PMSF);^38-40^ ligand **5** diisopropyl can phosphorylate the Ser329 residue via an O−P bond;^41^ ligand **6** is a substrate-mimic haloketone specifically designed for **PK**.^42^ Upon releasing the halogen atom (Cl), it forms a C−N bond between methylene and imidazole group in His174 residue, and a C−O bond between ketone carbonyl carbon and Ser329 residue. The complex structure between **PK** and ligands **6** has been deposited in PDB database (PDB entry: 4ZAR) but not published, the density map contains notably negative densities near His174 residue promoting further improvements.

We aim to apply the above workflow (Figure 1) to elucidate protein-ligand binding in proteinase K. When **PK** is cocktail soaked with ligands **4**-**6**, the resulting **PK**-**4, PK**-**5**, and **PK**-**6** complexes irreversibly formed and co-existed in the mixture (Figure S2B in Supporting Information). Ratio of each component is determined by binding affinities (K_D_) and ligand concentrations. In set E, proteinase K crystals were soaked for 12 hours in a cocktail of ligands **4**-**6** in equal concentrations (Table 1). The extended incubation time was determined from pilot experiments, ensuring a thorough reaction between ligands and **PK**. Three individual soaking sets F-H were set as references (Table 1). A total of 103 datasets were collected using automated MicroED (Figure 1B). The 82 datasets retained for further analysis (Figure S5 in Supporting Information; see “Method” for details) yielded 62 structures by molecular replacement without merging.^43^ The three ligands were unambiguously identified in the Fo-Fc maps after refinement (Figures 3 and 4C).

We compared the Fo-Fc densities in the binding site in cocktail set E with those from the crystals where a single ligand was individually soaked (reference sets F-H Figure 3). Overall, the densities are clearer when soaking with a single ligand compared to the cocktail soaks, albeit at a similarly weaker counter level (Figure 3A). For example, the phenyl group of ligand **4** is partially missing in the cocktail soak, but its distinctive shape and orientation still allows unambiguous identification. The ligand in the cocktail-soaked structure refines to ∼0.5 Å all-atom r.m.s.d of its arrangement in the individually soaked sets; Figure S7A in Supporting Information). The density of ligand **5** was complete and consistent in both sets E and G (Figure 3B), and the structures refined from these datasets showed identical binding geometries (Figure S7B in Supporting Information). In set E, the density of ligand **6** was complete and superior to the unpublished PDB structure 4ZAR (RMSD: 0.23 Å). Two covalent bonds between Ser329 and His174 residues were clearly visible from Fo-Fc maps (Figure 3C).^42^ In set H, the density of ligand **6** was also complete, but the His174 exhibited ∼2.9 Å shifts compared to set E; the density for Cl atom was visible. A non-covalent binding pose was occasionally observed in this case, where only non-covalent interactions were found between His174 and methylene (4.3 Å) and Ser329 and ketone carbonyl (3.0 Å). This alternative binding geometry might be caused by pH differences, which slowed the reaction rate of ligand **6** on **PK**.

Quantifying of each complex type in the mixture allowed for a quantitative comparison of their binding affinities: ligand 6 binds stronger than ligand **5**, which in turn exhibits greater affinity than ligand **4** (Figure 4C). Since the three ligands were mixed in approximately equal concentrations, the most frequently observed ligand **6** exhibited the highest binding affinity among the three. Ligand **5** showed slightly higher affinity than ligand **4**, but their counts are close, indicating comparable K_D_ values. MST measurements of K_D_ constants for ligands **4**-**6** were 260 µM for ligand **4**, N.A. for ligand **5**, and 18 µM for ligand **6**. Ligand **5** showed high activity in an aqueous environment and was degraded before a successful MST measurement; however, the measured K_D_ values for ligands **4** and **6** were consistent with MicroED results (Figures 4C-D).

## 3 Discussion

Cocktail soaking combined with high-throughput data collection can significantly accelerate drug discovery by allowing rapid crystallographic screening of ligands against target proteins.^1-6,9,10^ In conventional SC-XRD, which requires crystals at least ∼5 µm in size,^14^ the expected success rate of ligand binding is relatively low (∼1%) making it unsuitable for protein microcrystals smaller than this size threshold and also leads to significant waste of ligands. Weak-binding ligands might remain undetected if they failed in competition with stronger ligands in the cocktail, or untested if limited datasets were collected.

In this study, we proposed a workflow integrating cocktail soaking and automated MicroED for crystallographic ligand screening and structural analysis of protein-ligand complexes directly from microcrystals (Figure 1). After cocktail soaking, the mixture contains multiple protein-ligand complexes, all the ligands can be determined after sampling a sufficient number of crystals. Their Fo-Fc densities were complete and comparable to those in literature^26-28^ and reference sets (Figures 2 and 3). The ligand with the highest affinities is expected to form the most crystals after cocktail soaking. Quantifying the proportion of each complex allows the estimation of their relative binding affinities, which is consistent with literature^29,30^ and MST measured of K_D_ constants (Figure 4).

Detailed structural analysis of thermolysin and proteinase K in complex with their respective ligands reveals their binding differences in the active sites. For **TLN**, the phosphoryl group of ligand **1** coordinated with Zn ion at ∼2 Å, anchoring it within the binding cleft. The rest of molecule involved in at least 7 hydrogen bonds, 2 salt bridges, and 1 water bridge. Similarly, the sulfanyl group of ligand **2** coordinated with the Zn ion at ∼2.3 Å, forming 4 hydrogen bonds, 1 salt bridge, and 1 T-shaped *pi*-staking. There is no conserved binding pose observed for ligand **3**, and its orientation appeared flexible within the binding pocket, which can be temporarily stabilized by a hydrogen bond with Asn344 or Arg435 or a water molecule. As for **PK**, ligand **4** formed an O−S covalent bond with the Ser329 residue. Its sulfonyl group formed in at least 3 hydrogen bonds within the pocket. The rest of molecule (*e*.*g*. 2-aminoethyl benzyl group) might form a weak hydrogen bond with Gly239 residue (∼3.6 to 3.8 Å) with minimum structural constraints. This increased rotational freedom likely explained the incomplete densities observed in Fo-Fc maps (Figure 3A). Ligand **5** formed an O−P covalent bond with Ser329 residue, the phosphoryl group involved in 4 hydrogen bonds anchoring it within the binding pocket. As mentioned, ligand **6** displayed dual binding poses in sets E and H. In set E, ligand **6** formed 2 covalent bonds between His174 and methylene, Ser329 and ketone carbonyl carbon, which mimics the transition state of **PK** bound with substrate. It blocked the entry tunnel via at least 8 hydrogen bonds and 1 water bridge. In set H, ligand **6** was non-covalently interacted with at least 10 hydrogen bonds similar to set E.

The primary aim of this study is to develop a rapid crystallographic ligand screening approach, allowing the application of cocktail soaking into microcrystals. Because automated MicroED requires only 7 min per crystal, faster than a typical SC-XRD experiment conducted at synchrotron.^44^ More datasets can be collected in the same time. High-throughput data collection allows data collection to focus on high-quality datasets from crystals where ligands may be bound with high occupancy. Those datasets were subsequently trimmed by strict criteria (See “Method” for details) prioritizing crystals with better occupancy over high resolution. Most of the trimmed datasets enabled for unambiguous structural identification and refinement, however, cases where the same crystal binds several different ligands may be addressed using PanDDA in an “event map” approach,^45^ or as demonstrated by substituting with a better crystal with higher occupancy from the high-throughput datasets. Weakly diffracting crystals were excluded to leave only high-quality data solved for unambiguous structural determination (Figures 2 and 3).

This workflow offers future opportunities for drug discovery and development, such as the identification of novel drugs with superior binding compared to the biogenic ligands or existing drugs. It offers an alternative approach for evaluating ligand binding affinities, particularly for unstable compound like ligand **5**, for which directly measuring its K_D_ values is challenging (Figure 4C-D). Additionally, this method can yield a structure from each crystal, enabling characterization of the dynamics of protein-ligand interactions when many datasets are available (Figure 3C). In the follow-up experiments, more ligands can be introduced at the cost of requiring longer data collection time. Due to resolution limits, ligands selected for cocktail soaking must exhibit diverse shapes following the established literature practices.^1-6,9,10^ More structurally similar ligands can be tested in the future, given the resolution enhancement using crystal fragmentation^34^ or focused-ion-beam (FIB) milling.^46,47^ Lastly, multiple mutated protein crystals can be combined and soaked with ligands to investigate essential molecular contacts.

## 4 Methods

### 4.1 Materials

Thermolysin (*Bacillus thermoproteolyticus rokko*) was purchased from MedChemExpress. Proteinase K (*Tritirachium album*) was purchased from Sigma Aldrich. Ligands **1, 3, 6** were purchased from Sigma Aldrich; ligand **2** was purchased from MedChemExpress. Ligand **4** was purchased from Focus Biomolecules. Ligand **5** was purchased from Thermo Scientific Chemicals.

### 4.2 Protein crystallization

Crystallization of thermolysin and proteinase K followed previously reported procedures.^34,43^ Thermolysin was dissolved at a concentration of 80 mg/mL in 45% dimethyl sulfoxide, 50 mM Tris-HCl (pH 7.5), and 2.5 M cesium chloride. Sitting drops were set up by mixing 0.5 µL protein solution and 1.5 µL water and equilibrating over 300 µL water at room temperature. Proteinase K was dissolved at a concentration of 25 mg/mL in 50 mM Tris-HCl (pH 8.0) and mixed with an equal volume of 1.25 M ammonium sulfate (pH 6.5) at room temperature. Microcrystals formed overnight, see Figure S1 in Supporting Information.

### 4.3 Grid preparation

Crystal slurries of thermolysin and proteinase K were diluted and fragmented by sonicating for 20 s in an FS60 ultrasonic cleaner (Fisher Scientific) with a cold-water bath.^34^ The resulting crystal slurries were soaked with ligands in different sets (Table 1).

Holey carbon grids (Quantifoil, R2/2 Cu200) were glow-discharged for 30 s at 15 mA in negative mode. Blotting and vitrification were conducted using a Leica GP2 plunge freezer set at 20 °C and 90% humidity (Figure 1A). For each grid, 3 µL of crystal slurry were deposited on the carbon side and were gently blotted for 30 s from the copper side. Grids A1-H (corresponding to sets A1-H, Table 1) were plunged into liquid ethane and transferred to liquid nitrogen for storage.

### 4.4 Automatic MicroED data collection

MicroED data were collected with SerialEM on a Falcon 4i camera (4096 × 4096 pixels) mounted in a 300 keV Titan Krios Cryo-TEM (Thermo Fisher) equipped with a Selectris energy filter.^24,25,46,47^ Screening and data collection was accomplished using *in-house* SerialEM scripts: first, the atlas of whole grid was montaged under the imaging mode (LM 155×). Around 100 grid squares with proper blotting were selected and montaged at higher magnification (SA 2250×) taking ∼7.5 hours. Crystals with light contrast to the carbon film were typically thin and were chosen from montaged maps (∼1000 crystals).

The diffraction mode configured with a gun lens 8, spot size 11, 50 µm C2 aperture, 20 eV energy filter slit, 1.8 m or 2.5 m camera lengths.^46^ ∼10 µm sized parallel electron beam and a 100 µm SA aperture were used to result in a flux density of ∼0.002 e^-1^/(Å^2^·s) on samples.^46^ All the selected crystals were tested with 1 s exposure time at 0° tilt in the diffraction mode to assess resolution. Only the best 100 crystals with resolutions better than 3.5 Å (Figure 1B) were retained for data collection. Eucentric heights were automatically calibrated to maintain the crystals inside the SA aperture during the continuous rotation. A typical data collection used a constant rotation rate of ∼0.2°/s over an angular wedge of 100° from -50° to +50° in 420 s exposure time. Overnight automatic data collection was applied for grids A1, A2 and E (Table 1), which typically yielded ∼100 datasets per grid. Manual data collection was performed for grids B-D and F-H (Table 1) using the same settings, which typically resulted in 10-15 datasets per grid.

### 4.5 MicroED Data Processing

Electron-counted MicroED data were saved in EER format and converted to SMV format using mrc2smv (https://cryoem.ucla.edu/downloads).^47,48^ The converted frames were indexed and integrated in XDS,^49,50^ with the resolution cutoff chosen where I/sigma ≥1. Datasets with resolution <3.5 Å, completeness <75% were discarded. The remaining datasets were scaled in XSCALE^50^ and converted to MTZ format using XDSCONV.^50^ The structures were solved by molecular replacement using Phaser^51^ with electron scattering factors and unliganded PDB structures 5K7T and 6PKR as search models for thermolysin and proteinase K, respectively,^34,43^ and refined using Phenix.refine (Figures 2 and 3).^52,53^ Ligands were identified based on distinctive density shapes from initial Fo-Fc maps calculated without ligands, comparing to those from literature^26-28^ and individual soaking references (Figures 2 and 3). Ligands were finally added and refined to yield PDB structures for **TLN**-**1**/**2**/**3** and **PK**-**4**/**5**/**6** (Tables S1-S6 in Supporting Information).

### 4.6 Microscale thermophoresis (MST)

Labeled thermolysin was prepared in 25 mM HEPES (pH 7.4), 150 mM NaCl, labeled with RED-NHS dye (NanoTemper), and purified to a concentration of 5 mM. Ligands **1**-**3** were prepared in 25 mM HEPES (pH 7.4), 150 mM NaCl at a concentration of 2-5 mM, then diluted to a series of concentrations from mM to µM. The diluted ligands were mixed with labeled protein in 1:1 ratio, which were loaded to monolith capillaries (NanoTemper). MST was measured at 37 °C for 10 s with two repeats. MST measurements for proteinase K and ligands **4**-**6** followed the same procedure. DMSO was used in preparing 25 µM ligand **6** due to its low water solubility.

The normalized fluorescence signal (F_Norm_) versus the concentration of ligands [L] measured with MST are plotted in Figures 4B and 4D. The binding affinity (K_D_) was calculated by fitting F_Norm_ and [L] to the equation 1 below:

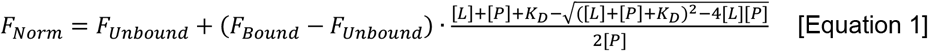

**where** *F*_*Norm*_is the normalized fluorescence signal, *F*_*Unbound*_, *F*_Norm_ is the signal of the target protein alone; *F*_*Bound*_: *F*_Norm_ signal of the complex. The final concentrations of the ligand and the target protein are denoted [*L*]: and [*P*]:, respectively; *K*_*D*_: is the binding affinity or dissociation constant.

## Acknowledgements

This study was supported by the National Institutes of Health P41GM136508. Portions of this research or manuscript completion were developed with funding from the Department of Defense MCDC-2202-002. Effort sponsored by the U.S. Government under Other Transaction number W15QKN-16-9-1002 between the MCDC, and the Government. The US Government is authorized to reproduce and distribute reprints for Governmental purposes, notwithstanding any copyright notation thereon. The views and conclusions contained herein are those of the authors and should not be interpreted as necessarily representing the official policies or endorsements, either expressed or implied, of the U.S. Government. The PAH shall flow down these requirements to its sub awardees, at all tiers. The Gonen laboratory is supported by funds from the Howard Hughes Medical Institute.

