## Supporting Information for "Ligand Screening and Discovery using Cocktail Soaking and Automated MicroED"

**Table S1**. Data processing and model refinement statistics for **TLN-1**.

| Name of the protein[a] | **TLN-1** | **TLN-1** | **TLN-1** |
|---|---|---|---|
| Set/Grid[b] | A1 | A2 | B |
| Accelerating Voltage (kV) | 300 | 300 | 300 |
| Wavelength (Å) | 0.0197 | 0.0197 | 0.0197 |
| Resolution range (Å) | 38.99 - 2.001<br>(2.08 - 2.0) | 44.44 - 2.599<br>(2.86 - 2.6) | 40.06 - 2.58<br>(2.84 - 2.58) |
| Space group | $P\ 6_1\ 2\ 2$ | $P\ 6_1\ 2\ 2$ | $P\ 6_1\ 2\ 2$ |
| Unit cell parameters (Å,°) | 94.66 94.66 132.96<br>90 90 120 | 94.32 94.32 132.77<br>90 90 120 | 92.51 92.51 128.72<br>90 90 120 |
| Total reflections | 290748 | 131162 | 140415 |
| Unique reflections | 24393 | 11334 | 10822 |
| Multiplicity | 11.92 | 11.57 | 12.97 |
| Completeness (%) | 99.7 | 99.9 | 99.9 |
| I/sigma (I) | 3.25 | 3.12 | 3.74 |
| R-meas | 0.576 | 0.616 | 0.583 |
| $CC_{1/2}$ | 0.943 | 0.927 | 0.969 |
| Wilson B-factor | 21.38 | 32.15 | 35.36 |
| Reflections used in refinement (#) | 24316 | 11285 | 10776 |
| Reflections used for R-free (#) | 1218 | 564 | 540 |
| R-work | 0.2565 | 0.2549 | 0.2100 |
| R-free | 0.2761 | 0.3114 | 0.2490 |
| Number of non-hydrogen atoms (#) | 2527 | 2489 | 2507 |
| macromolecules | 2469 | 2469 | 2469 |
| ligands | 5 | 5 | 5 |
| solvent | 53 | 15 | 33 |
| Protein residues (#) | 317 | 317 | 317 |
| RMS (bonds) | 0.004 | 0.002 | 0.002 |
| RMS (angles) | 0.63 | 0.44 | 0.50 |
| Ramachandran favored (%) | 96.50 | 95.86 | 95.22 |
| Ramachandran allowed (%) | 3.50 | 3.82 | 4.46 |
| Ramachandran outliers (%) | 0.00 | 0.32 | 0.32 |
| Rotamer outliers (%) | 1.19 | 0.40 | 0.00 |
| Clashscore | 2.51 | 2.72 | 3.77 |
| Average B-factor | 17.79 | 20.20 | 26.49 |
| macromolecules | 17.79 | 20.21 | 26.58 |
| ligands | 18.08 | 19.36 | 27.16 |
| solvent | 17.89 | 19.10 | 19.92 |

**Notes**: [a]Thermolysin was abbreviated as "**TLN**". [b]Data collected from experimental sets/grids A1-B, see Table 1.

**Table S2**. Data processing and model refinement statistics for **TLN-2**.

| Name of the protein[a] | **TLN-2** | **TLN-2** |
|---|---|---|
| Set/Grid[b] | A2 | C |
| Accelerating Voltage (kV) | 300 | 300 |
| Wavelength (Å) | 0.0197 | 0.0197 |
| Resolution range (Å) | 47.48 - 2.35 (2.53 - 2.35) | 44.54 - 2.77 (3.17 - 2.77) |
| Space group | $P\ 6_1\ 2\ 2$ | $P\ 6_1\ 2\ 2$ |
| Unit cell parameters (Å,°) | 94.96 94.96 133.45 90 90 120 | 94.44 94.44 134.22 90 90 120 |
| Total reflections | 179080 | 81213 |
| Unique reflections | 15416 | 9526 |
| Multiplicity | 11.62 | 8.53 |
| Completeness (%) | 99.7 | 99.6 |
| I/sigma (I) | 2.09 | 2.24 |
| R-meas | 0.755 | 0.660 |
| $CC_{1/2}$ | 0.911 | 0.900 |
| Wilson B-factor | 31.93 | 33.45 |
| Reflections used in refinement (#) | 15373 | 9491 |
| Reflections used for R-free (#) | 769 | 474 |
| R-work | 0.2731 | 0.2577 |
| R-free | 0.3014 | 0.2984 |
| Number of non-hydrogen atoms (#) | 2484 | 2468 |
| macromolecules | 2432 | 2432 |
| ligands | 22 | 22 |
| solvent | 30 | 14 |
| Protein residues (#) | 316 | 316 |
| RMS (bonds) | 0.003 | 0.002 |
| RMS (angles) | 0.49 | 0.47 |
| Ramachandran favored (%) | 96.18 | 96.18 |
| Ramachandran allowed (%) | 3.82 | 3.82 |
| Ramachandran outliers (%) | 0.00 | 0.00 |
| Rotamer outliers (%) | 0.40 | 1.59 |
| Clashscore | 2.53 | 1.90 |
| Average B-factor | 26.75 | 18.52 |
| macromolecules | 26.76 | 18.47 |
| ligands | 27.87 | 27.27 |
| solvent | 25.45 | 13.12 |

**Notes**: [a]Thermolysin was abbreviated as “**TLN**”. [b]Data collected from experimental sets/grids A2 and C, see Table 1.

**Table S3**. Data processing and model refinement statistics for **TLN-3**.

| Name of the protein[a] | **TLN-3** | **TLN-3** |
|---|---|---|
| Set/Grid[b] | A2 | D |
| Accelerating Voltage (kV) | 300 | 300 |
| Wavelength (Å) | 0.0197 | 0.0197 |
| Resolution range (Å) | 39.24 - 2.896 (3.32 - 2.9) | 47.34 - 2.23 (2.37 - 2.23) |
| Space group | P $6_1$ 2 2 | P $6_1$ 2 2 |
| Unit cell parameters (Å,°) | 94.7 94.7 135.03 90 90 120 | 94.68 94.68 131.66 90 90 120 |
| Total reflections | 87163 | 236138 |
| Unique reflections | 7131 | 17629 |
| Multiplicity | 12.22 | 13.39 |
| Completeness (%) | 83.8 | 99.7 |
| I/sigma (I) | 2.39 | 2.16 |
| R-meas | 0.666 | 0.733 |
| $CC_{1/2}$ | 0.887 | 0.940 |
| Wilson B-factor | 44.43 | 36.08 |
| Reflections used in refinement (#) | 7112 | 17580 |
| Reflections used for R-free (#) | 356 | 881 |
| R-work | 0.2890 | 0.2628 |
| R-free | 0.3298 | 0.2760 |
| Number of non-hydrogen atoms (#) | 2453 | 2494 |
| macromolecules | 2432 | 2432 |
| ligands | 12 | 12 |
| solvent | 9 | 50 |
| Protein residues (#) | 316 | 316 |
| RMS (bonds) | 0.002 | 0.003 |
| RMS (angles) | 0.43 | 0.51 |
| Ramachandran favored (%) | 93.95 | 95.86 |
| Ramachandran allowed (%) | 5.73 | 4.14 |
| Ramachandran outliers (%) | 0.32 | 0.00 |
| Rotamer outliers (%) | 0.79 | 1.59 |
| Clashscore | 2.75 | 2.97 |
| Average B-factor | 27.13 | 32.68 |
| macromolecules | 27.16 | 32.81 |
| ligands | 31.41 | 41.53 |
| solvent | 15.03 | 24.57 |

**Notes**: [a]Thermolysin was abbreviated as "**TLN**". [b]Data collected from experimental sets/grids A2 and D, see Table 1.

**Table S4**. Data processing and model refinement statistics for **PK-4**

| Name of the protein[a] | **PK-4** | **PK-4** |
|---|---|---|
| Set/Grid[b] | E | F |
| Accelerating Voltage (kV) | 300 | 300 |
| Wavelength (Å) | 0.0197 | 0.0197 |
| Resolution range (Å) | 43.27 - 2.02 (2.18 - 2.02) | 47.72 - 2.25 (2.48 - 2.25) |
| Space group | $P\ 4_3\ 2_1\ 2$ | $P\ 4_3\ 2_1\ 2$ |
| Unit cell parameters (Å,°) | 67.42 67.42 103.12 90 90 90 | 67.49 67.49 105.89 90 90 90 |
| Total reflections | 128645 | 112273 |
| Unique reflections | 14035 | 12043 |
| Multiplicity | 9.17 | 9.32 |
| Completeness (%) | 86.2 | 98.5 |
| I/sigma (I) | 3.72 | 4.56 |
| R-meas | 0.457 | 0.488 |
| $CC_{1/2}$ | 0.955 | 0.974 |
| Wilson B-factor | 23.63 | 32.09 |
| Reflections used in refinement (#) | 13959 | 12022 |
| Reflections used for R-free (#) | 698 | 601 |
| R-work | 0.2601 | 0.2127 |
| R-free | 0.3146 | 0.2721 |
| Number of non-hydrogen atoms (#) | 2075 | 2069 |
| macromolecules | 2029 | 2029 |
| ligands | 18 | 14 |
| solvent | 28 | 26 |
| Protein residues (#) | 279 | 279 |
| RMS (bonds) | 0.002 | 0.003 |
| RMS (angles) | 0.45 | 0.52 |
| Ramachandran favored (%) | 97.11 | 96.03 |
| Ramachandran allowed (%) | 2.89 | 3.97 |
| Ramachandran outliers (%) | 0.00 | 0.00 |
| Rotamer outliers (%) | 0.00 | 0.94 |
| Clashscore | 3.01 | 4.27 |
| Average B-factor | 18.40 | 26.11 |
| macromolecules | 18.34 | 26.03 |
| ligands | 25.72 | 41.87 |
| solvent | 18.47 | 23.54 |

**Notes**: [a]Proteinase K was abbreviated as "**PK**". [b]Data collected from experimental sets/grids E and F, see Table 1.

**Table S5**. Data processing and model refinement statistics for **PK-5**

| Name of the protein[a] | **PK-5** | **PK-5** |
|---|---|---|
| Set/Grid[b] | E | G |
| Accelerating Voltage (kV) | 300 | 300 |
| Wavelength (Å) | 0.0197 | 0.0197 |
| Resolution range (Å) | 40.74 - 1.97 (2.09 - 1.97) | 47.55 - 1.97 (2.09 - 1.97) |
| Space group | P $4_3$ $2_1$ 2 | P $4_3$ $2_1$ 2 |
| Unit cell parameters (Å,°) | 67.51 67.51 102.17 90 90 90 | 67.25 67.25 101.31 90 90 90 |
| Total reflections | 137334 | 139116 |
| Unique reflections | 17208 | 17111 |
| Multiplicity | 7.98 | 8.13 |
| Completeness (%) | 98.9 | 99.9 |
| I/sigma (I) | 3.35 | 4.23 |
| R-meas | 0.518 | 0.460 |
| $CC_{1/2}$ | 0.962 | 0.978 |
| Wilson B-factor | 22.65 | 28.98 |
| Reflections used in refinement (#) | 17174 | 17042 |
| Reflections used for R-free (#) | 859 | 851 |
| R-work | 0.2341 | 0.2016 |
| R-free | 0.2838 | 0.2315 |
| Number of non-hydrogen atoms (#) | 2082 | 2111 |
| macromolecules | 2029 | 2029 |
| ligands | 16 | 16 |
| solvent | 37 | 66 |
| Protein residues (#) | 279 | 279 |
| RMS (bonds) | 0.007 | 0.006 |
| RMS (angles) | 0.80 | 0.78 |
| Ramachandran favored (%) | 97.83 | 97.11 |
| Ramachandran allowed (%) | 2.17 | 2.89 |
| Ramachandran outliers (%) | 0.00 | 0.00 |
| Rotamer outliers (%) | 0.00 | 0.00 |
| Clashscore | 7.02 | 4.51 |
| Average B-factor | 18.90 | 24.93 |
| macromolecules | 18.85 | 24.87 |
| ligands | 26.57 | 34.36 |
| solvent | 18.17 | 24.21 |

**Notes**: [a]Proteinase K was abbreviated as "**PK**". [b]Data collected from experimental sets/grids E and G, see Table 1.

**Table S6**. Data processing and model refinement statistics for **PK-6**

| Name of the protein[a] | **PK-6** | **PK-6** |
|---|---|---|
| Set/Grid[b] | E | H |
| Accelerating Voltage (kV) | 300 | 300 |
| Wavelength (Å) | 0.0197 | 0.0197 |
| Resolution range (Å) | 34.7 - 1.801 (1.88 - 1.8) | 43.33 - 2.3 (2.53 - 2.3) |
| Space group | P $4_3$ $2_1$ 2 | P $4_3$ $2_1$ 2 |
| Unit cell parameters (Å,°) | 67.46 67.46 101.12 90 90 90 | 67.68 67.68 102.11 90 90 90 |
| Total reflections | 177311 | 99068 |
| Unique reflections | 22253 | 11100 |
| Multiplicity | 7.97 | 8.93 |
| Completeness (%) | 99.6 | 99.6 |
| I/sigma (I) | 3.95 | 3.25 |
| R-meas | 0.413 | 0.531 |
| $CC_{1/2}$ | 0.974 | 0.932 |
| Wilson B-factor | 20.1 | 27.2 |
| Reflections used in refinement (#) | 22226 | 11050 |
| Reflections used for R-free (#) | 1112 | 555 |
| R-work | 0.2269 | 0.2463 |
| R-free | 0.2574 | 0.2943 |
| Number of non-hydrogen atoms (#) | 2147 | 2101 |
| macromolecules | 2029 | 2029 |
| ligands | 43 | 44 |
| solvent | 75 | 28 |
| Protein residues (#) | 279 | 279 |
| RMS (bonds) | 0.006 | 0.002 |
| RMS (angles) | 0.80 | 0.46 |
| Ramachandran favored (%) | 97.83 | 94.95 |
| Ramachandran allowed (%) | 1.81 | 5.05 |
| Ramachandran outliers (%) | 0.36 | 0.00 |
| Rotamer outliers (%) | 0.94 | 0.00 |
| Clashscore | 5.49 | 2.25 |
| Average B-factor | 17.91 | 20.00 |
| macromolecules | 17.78 | 19.87 |
| ligands | 24.70 | 26.46 |
| solvent | 17.55 | 19.57 |

**Notes**: [a]Proteinase K was abbreviated as “**PK**”. [b]Data collected from experimental sets/grids E and H, see Table 1.

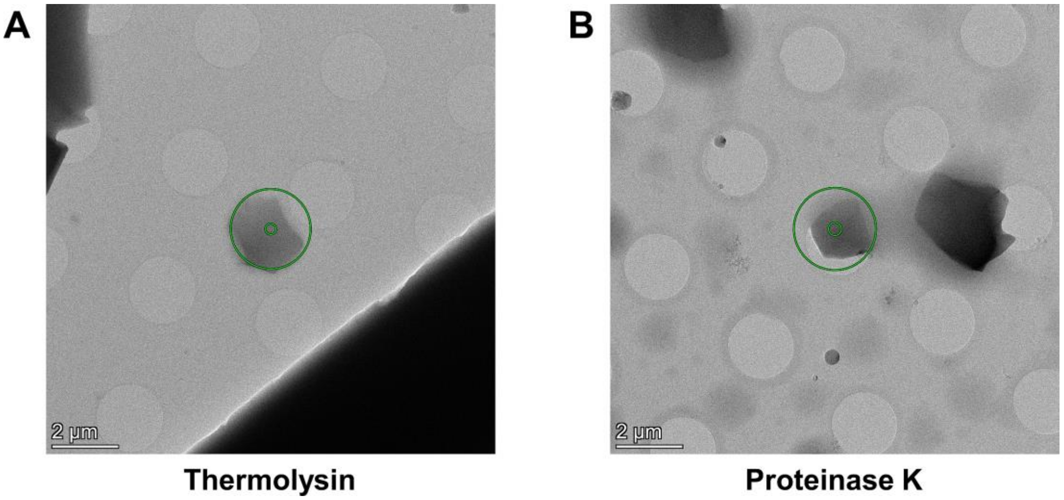


**Figure S1**. Representative images of microcrystals of (A) thermolysin and (B) proteinase K under imaging mode (2250 X).

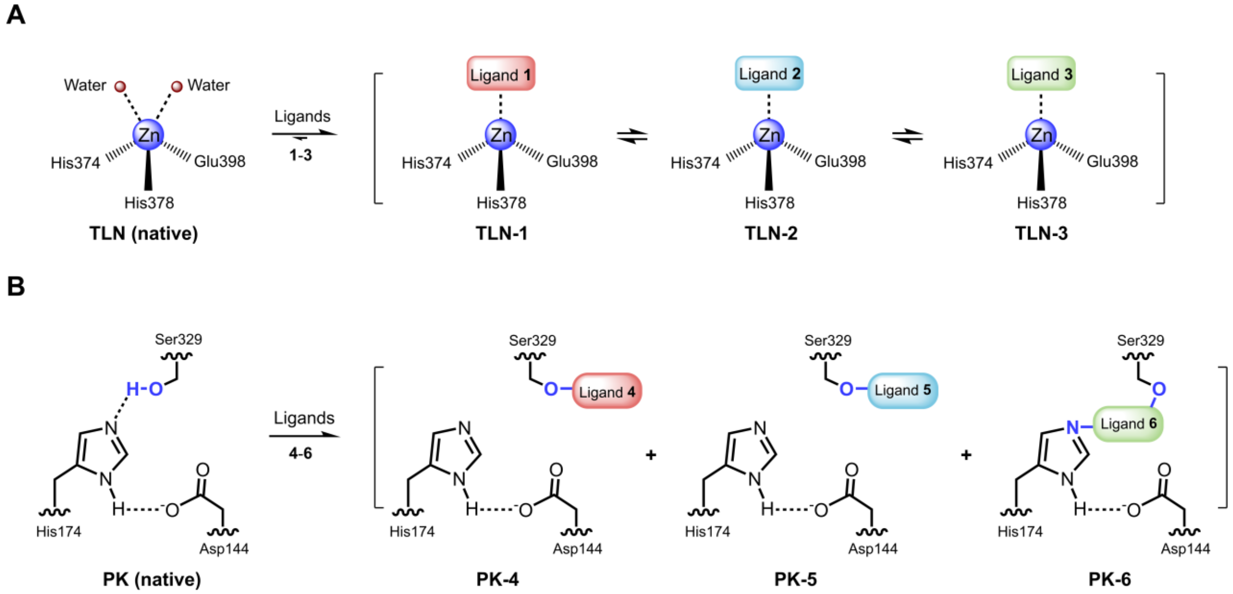


**Figure S2**. Schemes of protein-ligand binding mechanisms in cocktail soaking of (A) thermolysin with ligands **1**-**3** and (B) proteinase K with ligands **4**-**6**. Thermolysin was abbreviated as "**TLN**", proteinase K was abbreviated as "**PK**".

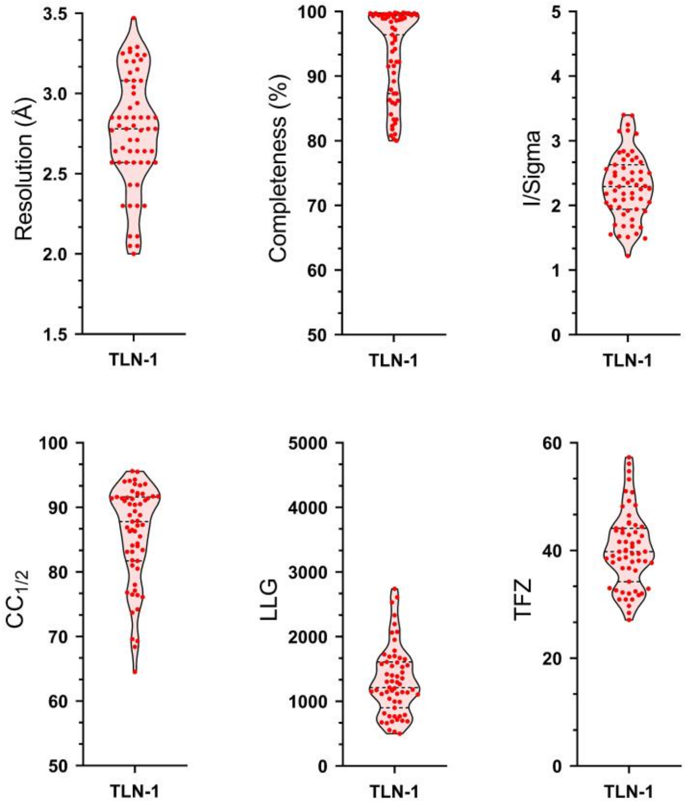


**Figure S3**. Statistics of **TLN-1** determined from cocktail soaking set A1. Thermolysin was abbreviated as “**TLN**”.

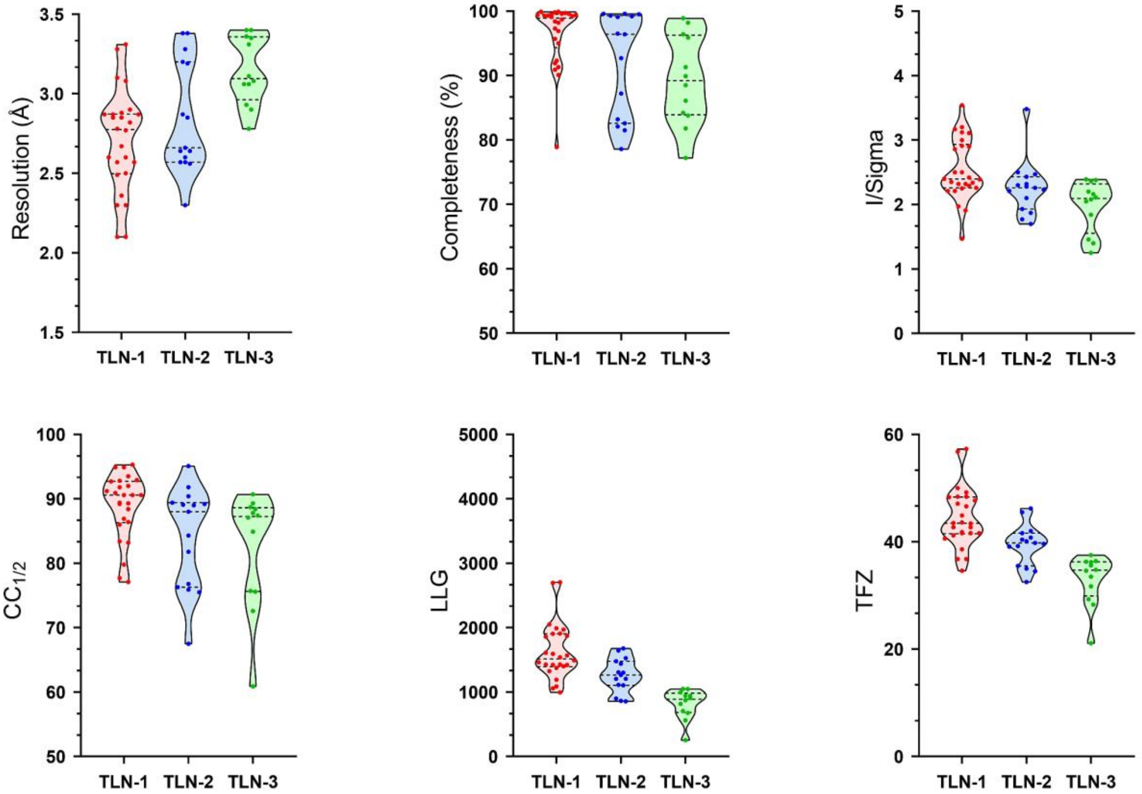


**Figure S4**. Statistics of **TLN**-**1**/**2**/**3** determined from cocktail soaking set A2. Thermolysin was abbreviated as "**TLN**".

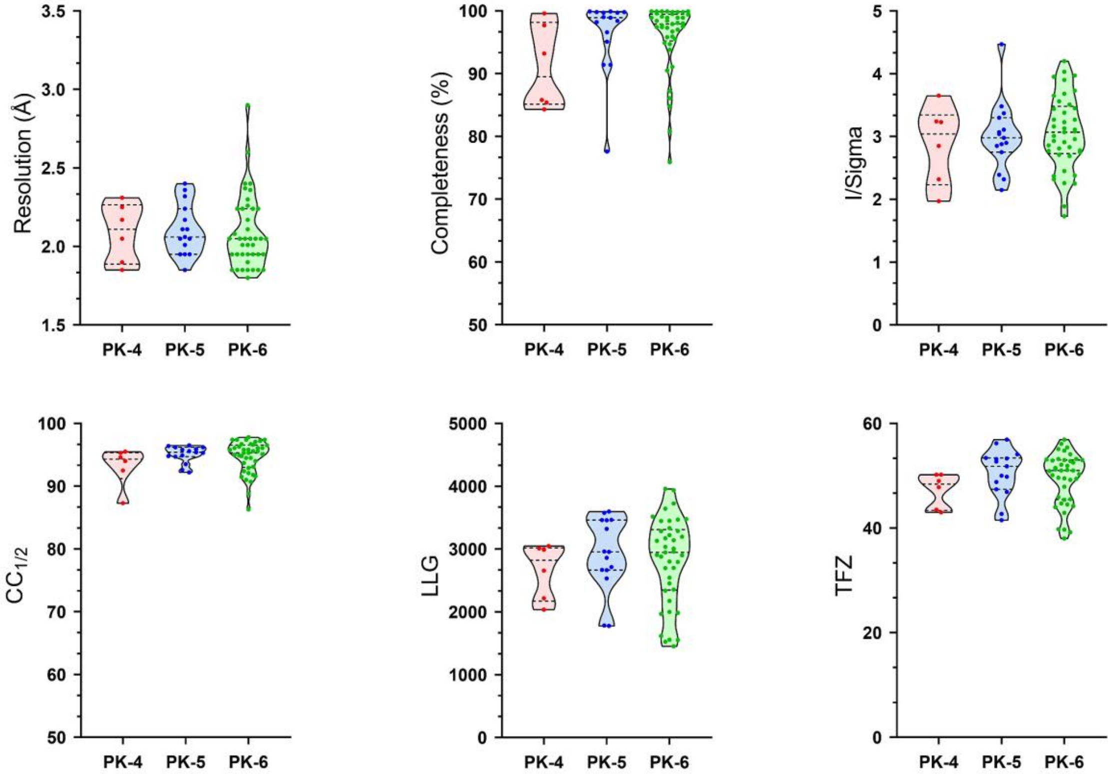


**Figure S5**. Statistics of **PK-4**/**5**/**6** determined from cocktail soaking set E. Proteinase K was abbreviated as “**PK**”.

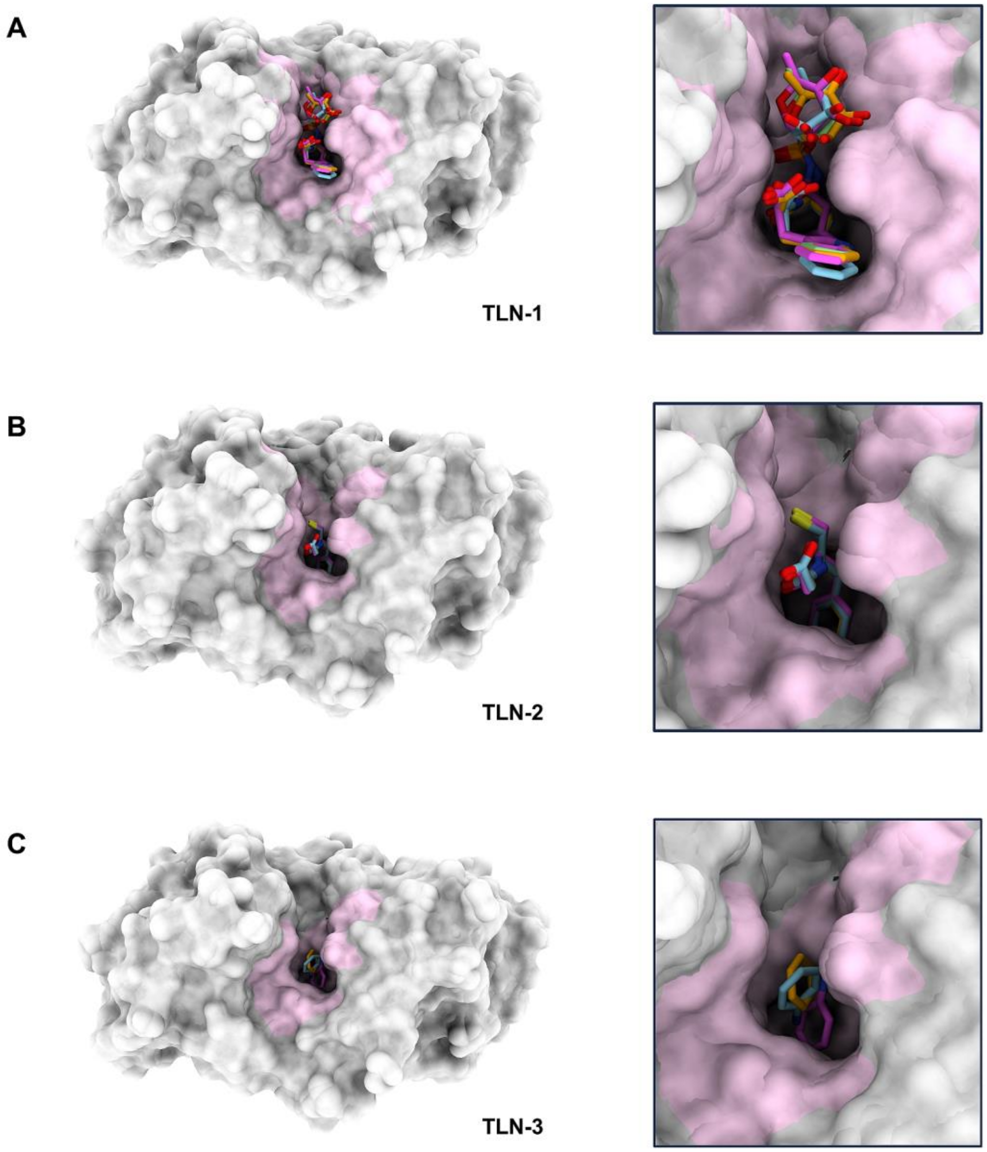


**Figure S6**. Overlay of **TLN-1**/**2**/**3** structures from different soaking sets and literature.[1-3] (A) **TLN-1** were colored by different sets: set A1, in orange; set A2, in blue; set B, in green; 1TLP, in magenta. (B) **TLN-2** were colored by different sets: set A2, in orange; set C, in blue; 1Z9G, in magenta. (C) **TLN-3** were colored by different sets: set A2, in orange; set D, in blue; 3MS3, in magenta. See Table 1 for details. Thermolysin was abbreviated as "**TLN**".

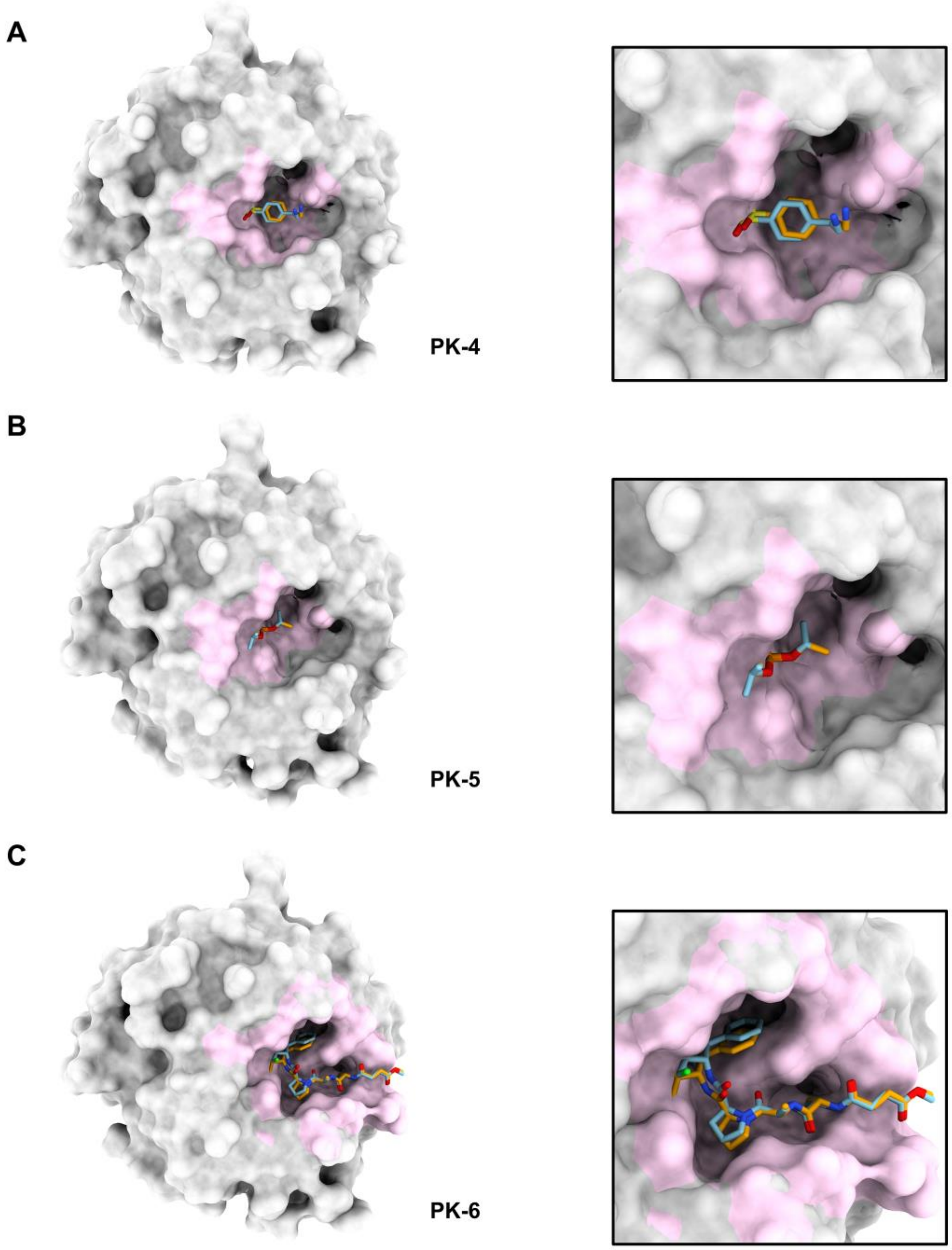


**Figure S6**. Overlay of **PK-4**/**5**/**6** structures from different soaking sets. (A) **PK-4** were colored by different sets: set E, in orange; set F, in blue. (B) **PK-5** were colored by different sets: set E, in orange; set G, in blue. (C) **PK-6** were colored by different sets: set E, in orange; set H, in blue. See Table 1 for details. proteinase K was abbreviated as “**PK**”.
